# Pregabalin silences oxaliplatin-activated sensory neurons to relieve cold allodynia

**DOI:** 10.1101/2022.06.20.496565

**Authors:** Federico Iseppon, Ana P. Luiz, John E. Linley, John N. Wood

## Abstract

Oxaliplatin is a platinum-based chemotherapeutic agent that causes cold and mechanical allodynia in up to 90% of patients. Silent NaV1.8-positive nociceptive cold sensors have been shown to be unmasked by oxaliplatin and other neuropathic insults. This event has been causally linked to the development of cold and mechanical allodynia. Pregabalin is an anti-epileptic and analgesic drug that acts through a calcium channel α_2_δ-1 subunit to lower neurotransmitter release. Recent data also suggest pregabalin can act on NMDA receptors and other proteins, but the site of analgesic action has been considered to be the central nervous system. We examined the effects of pregabalin on oxaliplatin-evoked unmasking of cold sensitive neurons using mice expressing GCaMP-3 driven by a *Pirt* promoter in all sensory neurons. We found that in mice treated with oxaliplatin, intravenous injection of pregabalin significantly decreased cold allodynia. Interestingly, pregabalin also decreased the number of sensory neurons responding to cold nociceptive stimuli by altering their excitability and their temperature thresholds. These silenced neurons are medium/large cells responding to both painful mechanical and cold stimuli, corresponding to the “silent” cold sensors that become active in numerous neuropathic pain models. Deletion of α_2_δ-1 subunits abolished the effects of pregabalin on both cold allodynia and the silencing of sensory neuron unmasked by oxaliplatin. Taken together, these results define a novel, peripheral inhibitory effect of pregabalin on the excitability of silent cold-sensing neurons in a model of oxaliplatin-dependent cold allodynia.

**Abbreviated Summary:** Iseppon et al. report a novel, peripheral effect of pregabalin on oxaliplatin-dependent cold allodynia. The drug exerts its effect by silencing a specific sub-population of neurons responding to cold and mechanical stimuli in the dorsal root ganglion, and this effect is dependent on the α_2_δ-1 subunit of voltage-gated calcium channels.

## Introduction

Cold allodynia is a debilitating condition where patients suffering from it experience mild, innocuous cooling as severe pain^1^. This ailment is common in many neuropathic pain conditions, from nerve injury to diabetic neuropathy, toxin poisoning and chemotherapy treatment, with an occurrence of up to 90% of patients^2^. The pathophysiological mechanisms of cold allodynia are currently unclear, but a recent article showed a common feature of many neuropathies carrying this condition: a previously unidentified population of large-diameter, silent cold-sensing neurons that become active and contribute to cold allodynia in these states^3^. This shift in neuronal activity was particularly evident in an intra-plantar pain model of oxaliplatin-induced neuropathy, one of the conditions that has the highest occurrence of peripheral cold allodynia^2–5^. Oxaliplatin is a chemotherapeutic agent associated with acute neurotoxicity that manifests as mechanical and cold allodynia, as well as chronic neurotoxicity on long-term treatment due to cumulative dosage^6^.

There is still no clear therapeutic route to alleviate neuropathic pain symptoms and cold allodynia. Few drugs have shown meaningful results, but gabapentinoids show a promising analgesic function in several neuropathic pain models^7–12^. These compounds are thought to act primarily on excitability through their interaction with Voltage-Gated Calcium Channels (VGCCs) via the auxiliary subunit α2δ-1^12,13^, although they can have other molecular targets such as NMDA receptors^12,14–16^. Gabapentin, and its successor pregabalin, are thought to act both at spinal and supra-spinal loci, inhibiting ascending nociceptive input from Dorsal Root Ganglion (DRG) neurons and activating descending inhibitor pathways to ameliorate neuropathic pain^12,15^.

To investigate the functional effects of pregabalin on an acute, oxaliplatin-dependent model of cold allodynia, we used behavioural testing as well as *in vivo* calcium imaging to dissect the effect of this compound on DRG neuronal activity. Here we describe a novel, peripheral action of pregabalin that results in the preferential silencing of the large-diameter, silent cold-sensing neurons involved in cold allodynia ^3^. Furthermore, we show that this effect is dependent on the α_2_δ-1 subunit of the VGCCs and is abolished in a global α_2_δ-1 transgenic knockout mouse line.

## Materials and methods

### Experimental Model and Treatment

#### Mice

All animal procedures carried out at University College London were approved by University College London ethical review committees and conformed to UK Home Office regulations (Animals (Scientific Procedures) Act 1986). For experiments using only wild-type mice, adult (>6 weeks-old) C57BL/6 mice from Charles River were used. Both transgenic mice used in this paper had a C57BL/6 background. *Pirt*-GCaMP-3 mice were kindly provided by Prof. Xinzhong Dong (John Hopkins University, Baltimore, MD, USA)^17^ and global α_2_δ-1 knock-out (KO) mice were generously donated by Prof. Annette Dolphin (University College London, London, UK)^18^.

All experiments using genetically altered lines were performed on adult male and female α_2_δ-1-KO mice and wild type littermates, obtained by breeding heterozygotes. Mice were housed in groups of no more than 5 on a 12 h:12 h light dark cycle; food and water were available *ad libitum*. Both male and female animals were used for all experiments, in equal numbers where possible. These studies were not however designed to test for sex differences, and sexes were pooled for analysis. The number of animals used to generate each dataset is described in individual figure legends. For genotyping, genomic DNA was isolated from ear tissue followed by PCR.

#### GCaMP Virus injection

Pups from α_2_δ-1 breeding heterozygote mice (P2 to P5) were used for injection of GCaMP expressing virus. Before the injection, the mother was anesthetized with a low concentration of isoflurane (1%) and pups were anesthetized by hypothermia. Virus pAAV.CAG.GCaMP6s.WPRE.SV40 (Plasmid #100844, Addgene) was injected into the plantar area of the hindpaw using a 10 μl Hamilton syringe with a cannula connected to a 30G needle. 5 μl of virus (titre = 5 × 10^12^ GC/ml) was injected into each of the two hindpaws. After injection, the pups were kept on a heating box until their body temperature returned to normal. Then they were returned into the cage before the mother was put back and rubbed against the pups. The mice injected with virus, after weaning, had the genotyping for α_2_δ-1 subunit checked by PCR. Both sexes were used for *in vivo* imaging 6 to 8 weeks after injection.

#### Chemotherapy-induced neuropathy pain model

Chemotherapy-induced neuropathy was studied in mice using the intraplantar oxaliplatin model first described by Deuis *et al* because this treatment recapitulates the rapid onset of cold allodynia in human patients infused with the drug^4^. Oxaliplatin (Sigma) was made up in 5% glucose solution in Milli-Q water to an equivalent dose of 80 μg in 40 μl. This is because oxaliplatin is unstable in chloride-containing saline solution. Mice were treated by intraplantar injection into the left hindpaw. Behavioural testing or imaging was assessed at least 3 hours after injection.

#### Treatment

After injection of oxaliplatin and the behavioural or calcium imaging baseline measurements (at least 3 hours after injection), the mice were treated with pregabalin (2 mg/kg) or vehicle (PBS solution) via intravenous (i.v.) injection. Behavioural testing or calcium imaging were performed up to 60 or 50 minutes after injection respectively.

### Behavioural Testing

All animal experiments were performed in accordance with Home Office Regulations. The investigator was blind to treatment and/or genotype. Animals were acclimatized to handling and every effort was made to minimize stress during the testing. Both male and female animals were used.

#### Cold Plate

Mice were placed on the Cold Plate apparatus (Ugo Basile®). The cold plate was maintained at 10°C for 5 minutes while the animal was free to move around on the plate and the number of nociceptive behaviours (shaking, lifting, licking, guarding, biting) displayed by the injected paw were counted by the observer ^19^.

### *In Vivo* Calcium Imaging

#### Acquisition

Adult mice expressing GCaMP-3 (6 to 12 weeks, male and female) were anesthetized using ketamine (100 mg/kg), xylazine (15 mg/kg) and acepromazine (2.5 mg/kg). Depth of anaesthesia was confirmed by pedal reflex and breathing rate. Animals were maintained at a constant body temperature of 37°C using a heated mat (VetTech). Lateral laminectomy was performed at spinal level L3-5. In brief, the skin was incised longitudinally, and the paravertebral muscles were cut to expose the vertebral column. Transverse and superior articular processes of the vertebra were removed using OmniDrill 35 (WPI) and microdissection scissors. To obtain a clear image of the sensory neuron cell bodies in the ipsilateral dorsal root ganglion (DRG), the dura mater and the arachnoid membranes were carefully opened using microdissection forceps. Artificial spinal fluid (values are in mM: 120 NaCl, 3 KCl, 1.1 CaCl_2_, 10 Glucose, 0.6 NaH_2_PO_4_, 0.8 MgSO_4_, 18 NaHCO_3_, pH 7.4 with NaOH) was constantly perfused over the exposed DRG during the procedure to maintain tissue integrity. The animal was mounted onto a custom-made clamp attached to the vertebral column, rostral to the laminectomy. The trunk of the animal was slightly elevated to minimize interference caused by respiration. The DRG was isolated by coating with silicone elastomer.

Images were acquired using a Leica SP8 confocal microscope. A 10x dry, 0.4-N.A. objective with 2.2 mm working distanced was used, with image magnification of 0.75-3x optical zoom. GCaMP-3 was excited using a 488 nm laser line (3-10% laser power). GCaMP signal was detected using a hybrid detector (60% gain). 512 × 512 pixel images were captured at a frame rate of 1.55 Hz, bidirectional scan speed of 800 Hz, and pixel dwell time of 2.44 μs.

Noxious and innocuous stimuli were applied to the left hindpaw, ipsilateral to the exposed DRG. For thermal stimuli, the ventral side of the paw was immersed with ice-water (nominally 0°C), acetone (100%) or water heated to 55°C using a Pasteur pipette. For delivery of precise temperature stimuli, a Peltier-controlled thermode (Medoc) was used. For mechanical stimuli, a pinch with serrated forceps was used. An interval of at least 30 s separated each stimulus application.

#### Analysis

Image stacks were registered to a reference image – typically, the first frame in the series – using the FIJI plugin TurboReg (accurate rigid body transformation) to correct for XY drift. Stacks that showed excessive Z movement were excluded from analysis. Regions of interest (ROI) were manually drawn around apparently responding cells using the polygon tool in FIJI. Mean pixel intensity over time for each ROI was extracted and analysed. The time series of mean pixel intensity for each ROI was smoothened by a five time-point rolling average to remove high-frequency noise. Next, we calculated the derivative of the mean pixel intensity. We calculated a mean baseline derivative for the 20 seconds preceding stimulus application. Neurons were classed as responders if, within 30 s of stimulus application, the maximum derivative was greater than the baseline derivative plus four standard deviations – that is, a Z-score of at least 4. We then calculated the ΔF/F_0_ value for each response to obtain a normalized measure of change in fluorescence. Neurons which showed a ΔF/F_0_ less than 0.25 were then discarded. Each trace was then manually screened as a further precaution against false positives. The remaining neurons that made up the responding population were then used for statistical analysis.

### Quantification and Statistical Analysis

For *in vivo* imaging experiments, *n* refers to the number of cells responding to any stimulus. For all imaging and data, the number of animals used is indicated in the legend. For behavioural experiments, *n* refers to the number of animals. No power calculations were performed; however, sample sizes are similar to those used in the field.

Datasets are presented using appropriate summary statistics as indicated in the legend, typically accompanied by raw data points or a representation of the underlying distribution. Behavioural data were generally assumed to be normal, as is typical in the field, and error bars denote mean ± Standard Error of Measurements (SEM). For *in vivo* imaging experiments, cells from all animals were pooled for analysis. Normality was not assumed when comparing cross-sectional areas or response magnitude of responding cells. Data are summarized using averages and SEMs, as well as cumulative or numerical distribution plots.

Tests of statistical comparison for each dataset are described in detail in figure legends. When comparing two treatment timepoints, paired *t* test was used. When comparing the distribution of cell cross-sectional areas for two groups, the Wilcoxon test was used. For more than two groups, One-Way ANOVA or Kruskal-Wallis test were used with post-hoc tests corrected for multiple comparisons. When comparing the effect of two factors on multiple groups, a repeated-measures Two-Way ANOVA was used, with post-hoc tests corrected for multiple comparisons. Curve fitting was performed using linear regression or non-linear regression functions.

Statistical tests were all performed using GraphPad Prism 9. An α-value of *p<*0.05 for significance testing was used. All p-values resulting from planned hypothesis testing are reported.

## Results

### Pregabalin ameliorates oxaliplatin-dependent cold allodynia by suppressing silent cold-sensing neurons

#### Pregabalin alleviates cold allodynia in oxaliplatin-dependent neuropathy

Cold allodynia following oxaliplatin treatment is a side effect that occurs in humans within a few hours from the injections. The model used in this paper, developed to isolate the activity of oxaliplatin on peripheral sensory neurons, consists of an intraplantar injection of 80μg (in 40μl) of oxaliplatin that elicits strong nociceptive responses, that comprise licking, flinching, and shaking of the injected paw as soon as 3-4 hours from the injection, assessed using a cold plate assay^3,4^. We observed the same increase in nociceptive responses upon oxaliplatin injection when performing a cold plate assay over 5 minutes at 10° C (Supplementary Fig. 1). When the mice were treated with 2mg/kg of pregabalin via intravenous injection, the number of nociceptive behaviours decreased significantly from 105 (± 5.5) to 56 (± 4.1). Control mice injected with vehicle showed a slight decrease in the total number of behaviours with high variability and no significant difference from the untreated ones (87 ± 12.2) (Figure 1A).

**Figure 1.**
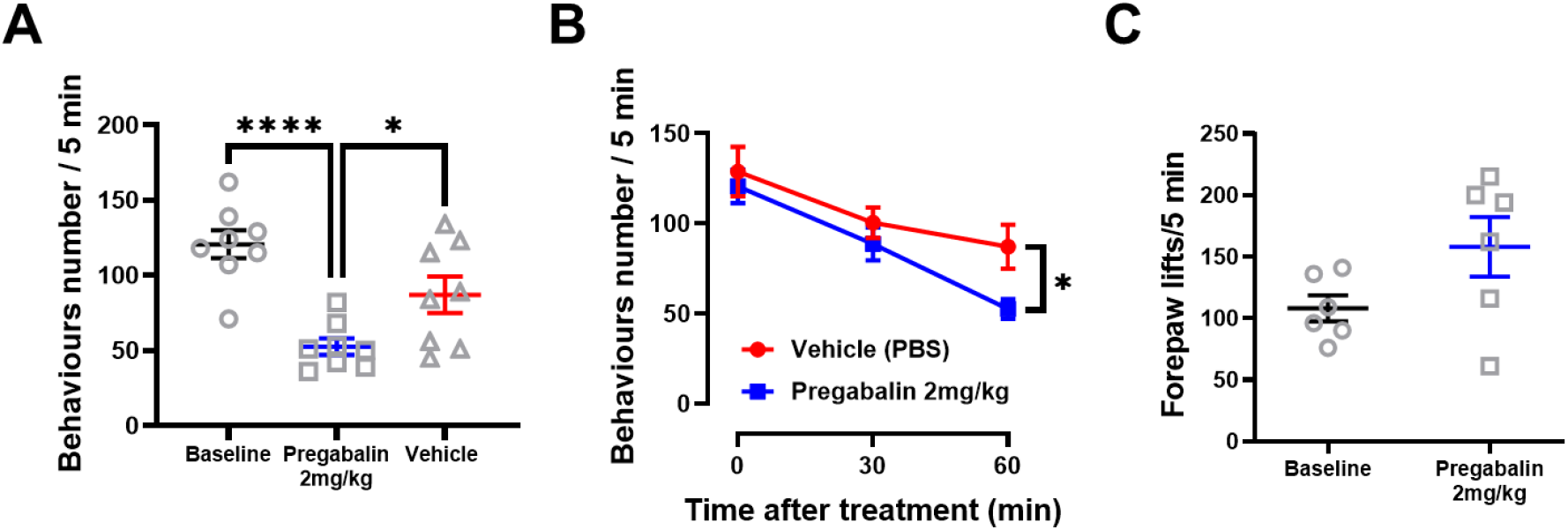
Assessment of the effect of pregabalin on acute oxaliplatin-induced cold allodynia. (A) Cold-plate assessment at 10°C of mice treated with oxaliplatin and successively either vehicle or 2 mg/kg of pregabalin. Activity was measured as the total number of nociceptive behaviours (hindpaw lifts, shakes, licks) over the test duration. A cutoff time of 300s was used to limit tissue damage. The baseline data in the graph refers to the mice treated with pregabalin. (B) Plot showing the decrease of the number of nociceptive behaviours over time following vehicle or pregabalin injection. (C) Cold-plate assessment at 10°C of naïve mice treated with pregabalin. Activity was measured as the total number of forepaw lifts over the test duration. A cutoff time of 300s was used to limit tissue damage. *n* = 8 oxaliplatin + pregabalin mice, *n* = 8 oxaliplatin + vehicle mice, *n* = 6 naïve + pregabalin mice Statistical analyses in (A) and (B) were performed using One-Way and Two-Way ANOVA tests with multiple comparisons respectively. Statistical Analysis in (C) was performed using paired t-test. **** = P<0.0001; * = P<0.05.

Nociceptive responses to cold temperature were measured 30 and 60 minutes after intraplantar injection, and a significant difference between the mice treated with pregabalin and those treated with vehicle is evident only after 60 minutes (Figure 1B). Next, we assessed the specificity of the effect of pregabalin on cold allodynia by treating mice with 2mg/kg pregabalin without the prior oxaliplatin injection. In this case no difference was observed in the behaviour of the mice maintained 5 minutes on a 10° C cold plate (Figure 1C).

#### Pregabalin silences cold-sensing cells by decreasing their excitability

Next, to dissect the effect of pregabalin on cold allodynia, we used *in vivo* calcium imaging to investigate if and how pregabalin treatment affects the responses of sensory neurons to various mechanical and thermal stimuli, with particular attention to any changes in their responses to cold temperatures. It is known that the acute oxaliplatin-dependent neuropathy model used in this study causes a change in the peripheral representation of cold, with the appearance of a population of large, normally silent, cold responding cells^3^. Using laser-scanning confocal imaging, we performed one-photon calcium imaging on the L4 DRG of oxaliplatin-treated mice to image the calcium signals of the neurons’ somata. Our experiments show a similar result, with a population of cells responding to ice-water and acetone stimuli of 27% and 17%, respectively (Figure 2A, B) (Supplementary Fig. 2). Interestingly, after intravenous injection of 2 mg/kg of pregabalin, we observed a decrease of the percentage of cells responding to either ice-water or acetone that becomes significant as early as 40 minutes after injection and decreases to 13% and 8% 50 minutes after injection respectively (Figure 2 B) (Supplementary Fig. 2B). Moreover, following pregabalin treatment, we observed a decrease in the intensity of the calcium signals evoked in cold-responding cells: both the peak fluorescence intensity (ΔF_max_) and the area under the curve decreased significantly after 40-50 minutes from the injection of pregabalin (Figure 2C-F).

**Figure 2.**
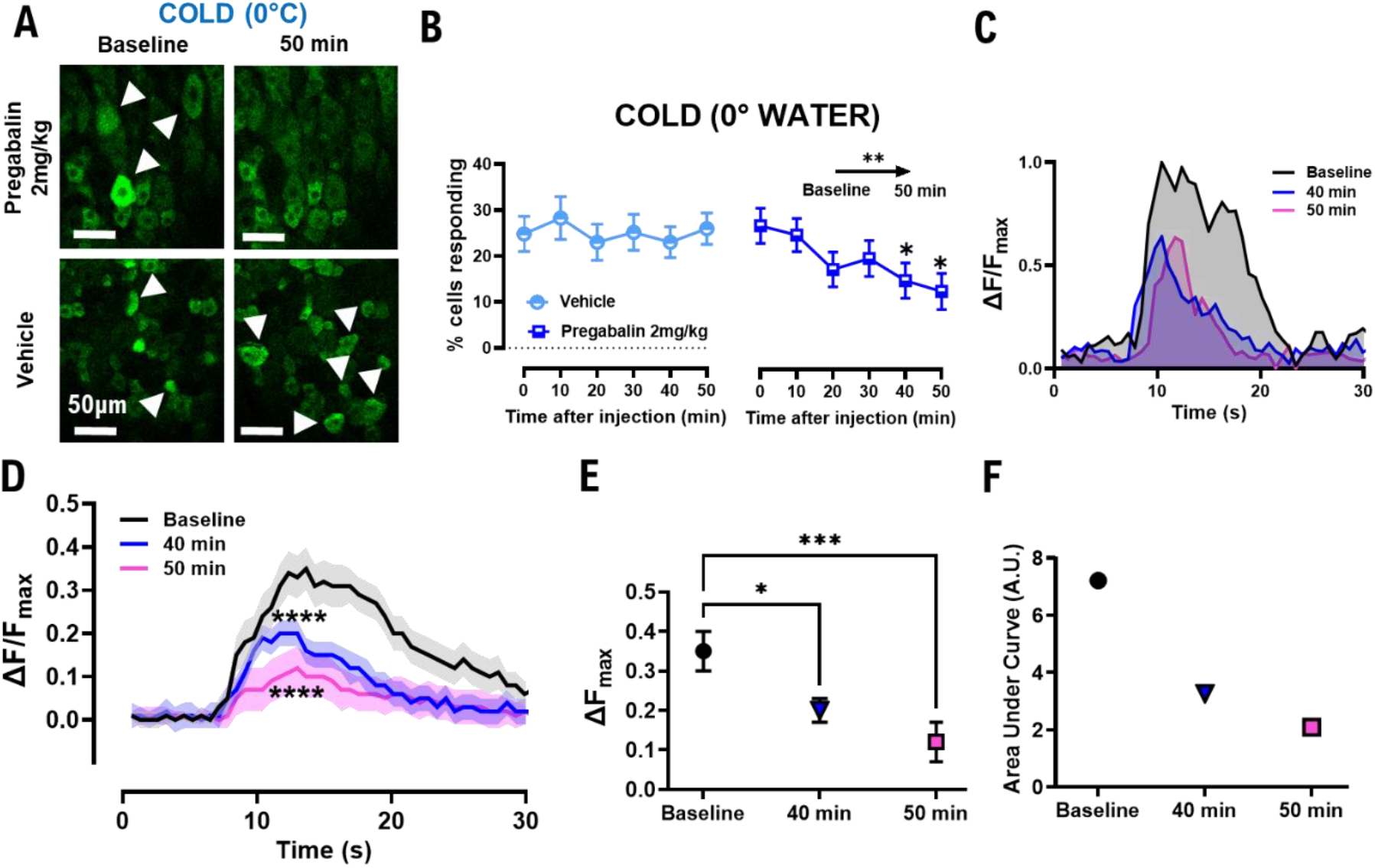
Pregabalin treatment decreases both the number of neurons responding to cold stimuli and the intensity of their responses. (A) Example images showing the reduction in the population of DRG neurons responding to ice-water 50 minutes after treatment with 2 mg/kg of pregabalin. (B) Graph showing the decrease of the percentages of cell responding to ice-water stimulus. (C) Example traces showing the decrease in the intensity of the calcium signals over time from pregabalin injection. (D) Plot showing the mean response amplitude before (black trace) and 40 (blue) and 50 (pink) minutes after pregabalin injection. *n* = 50 cells for all time points. (E) Graphs showing the ΔF_max_ changes before and 40 and 50 minutes after pregabalin injection. (F) Graphs showing the area under curve changes before and 40 and 50 minutes after pregabalin injection. *n* = 6 pregabalin-treated mice, *n* = 5 vehicle-treated mice. Statistical analysis in (B) was performed using Repeated Measures ANOVA test with multiple comparisons. Statistical analyses in (D) and (E) were performed using One-way ANOVA. * = P<0.05; ***=P<0.01; *** = P<0.001; ****=P<0.0001.

This effect of pregabalin is specific to cold sensory stimuli, the ones most significantly impacted by oxaliplatin: we observed no significant change in the number of cells responding to either mechanical or heat stimulation at any point during the treatment with pregabalin (Supplementary Fig. 3).

The oxaliplatin treatment is known not to affect the temperature threshold or excitability of cold sensing neurons, while unmasking silent cold nociceptors^3^. We investigated any possible change in the thermal activation thresholds of cold responding cells after treatment with pregabalin by delivering transient 4°C temperature drops, from 30 to 2°C, through a Peltier-controlled thermode (Figure 3A). While at baseline the cell recruitment rises in a linear fashion with temperature decrease, after treatment with pregabalin the linear correlation skews towards lower temperatures: this is a significant change and an indicator that pregabalin increases the activation threshold of cold responding cells (Figure 3B) (Supplementary Fig. 4).

**Figure 3.**
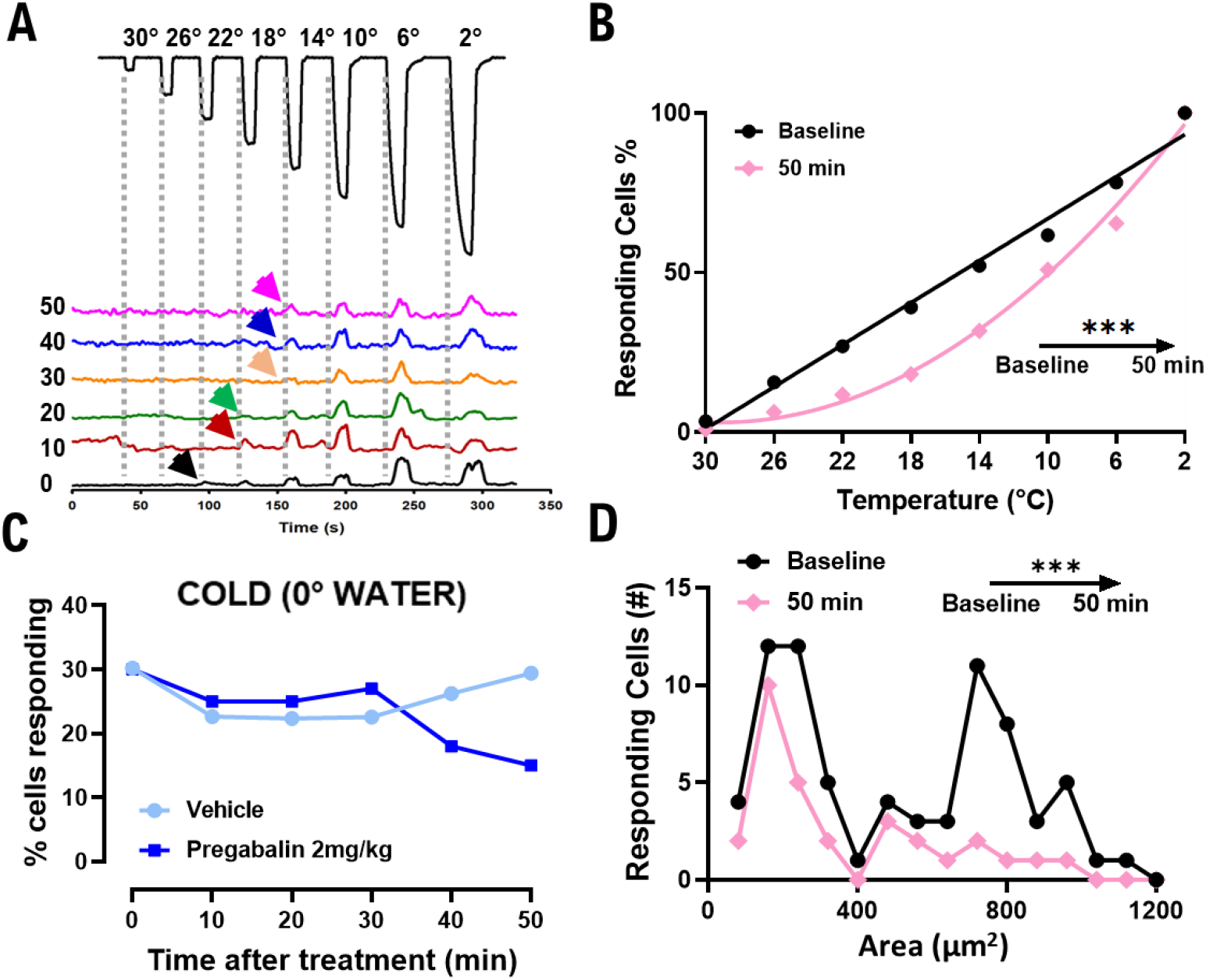
Pregabalin treatment significantly reduces the excitability of cold responding cells and targets preferentially medium-large mechano-cold sensors. (A) Example traces of a representative cold-sensing cell in response to a series of 4°C temperature drops from a holding temperature of 32°C. The traces represent the change in the threshold of this neuron over time (0-50 minutes) from pregabalin injection. (B) Graph showing the change of the relationship between the number of cold-responding neurons and the temperature drop over time from pregabalin injection. While the relationship at baseline can be fit with a linear equation (y = -3,292x + 99,84) (r^2^=0.9880) (*n* = 115), the one at 50 minutes fits a quadratic equation (y= 110,8 - 7.381x + 0,1265x^2^) (r^2^=0.9927) (*n* = 109). The slopes at baseline and 50 minutes after treatment are significantly different (P<0.001), as analyzed by Wilcoxon test. (C) Graphs showing the change in the number of total mechano-responding neurons that respond also to ice-water stimulus. Mice treated with vehicle do not exhibit the decrease over time from treatment as the mice treated with pregabalin do. (D) Numeric plots of the distribution of cold-responding cell areas in oxaliplatin-treated mice before and after 50 minutes from pregabalin injection. The difference between the distributions at different timepoints after treatment is statistically significant (P=0.0001), as analyzed by Wilcoxon test. *n = 7* mice for temperature threshold experiments; *n* = 6 pregabalin-treated mice, *n* = 5 vehicle-treated mice.

#### Pregabalin treatment suppresses predominantly ‘silent’ cold sensors

Acute oxaliplatin injection indeed causes the recruitment of cells that do not respond to cold stimuli before treatment. It also increases receptor polymodality, with a marked increase in neurons responding to both mechanical and cold stimuli: the population of silent mechano-sensors primarily respond to high-threshold mechanical stimuli^3^. We observed similar percentages in our model, and furthermore the injection of pregabalin decreased the population of cells responding to both mechanical and ice-water and acetone from 30% to 15% and from 20% to 6% respectively (Figure 3C) (Supplementary Fig. 5A). This effect is specific to mechano-cold responding cells, as the percentage of mechano-heat polymodal neurons did not vary significantly after treatment in either condition (Supplementary Fig. 5B). When analysing the area of cold-responding neurons we find the same two different populations found in MacDonald et al, 2021. Pregabalin treatment significantly affects the peripheral representation of cold allodynia: the analysis of the cross-sectional area of cold-responding cells shows a significant difference between the area of responding neurons after 50 minutes from pregabalin injection. Indeed, the treatment, although not completely, preferentially silences larger cold-responding cells, those ‘silent’ cold sensors that are unmasked in the oxaliplatin-dependent neuropathy (Figure 3D) (Supplementary Fig. 6).

Taken together, these results highlight the presence of a new, peripheral effect of pregabalin that inhibits the activity of de-novo cold sensing neurons by decreasing their excitability and thus ameliorates cold allodynia in a model of acute oxaliplatin-dependent neuropathy.

#### The peripheral effect of pregabalin is dependent on the Voltage Gated Calcium Channel subunit α_2_δ-1

Next, we investigated the mechanism of action of pregabalin on the silencing of the unmasked cold sensors by studying the role of its best-known target: α_2_δ-1. We performed the same cold plate assay described before on WT and global α_2_δ-1-KO mice: we treated all mice with oxaliplatin and 2 mg/kg of pregabalin, and we observed a significant difference between the two groups. Moreover, when the results were compared to the previous ones, there is an overlap between the vehicle-treated group and the α_2_δ-1-KO mice, suggesting an almost complete loss of the effect of pregabalin (Figure 4A).

**Figure 4.**
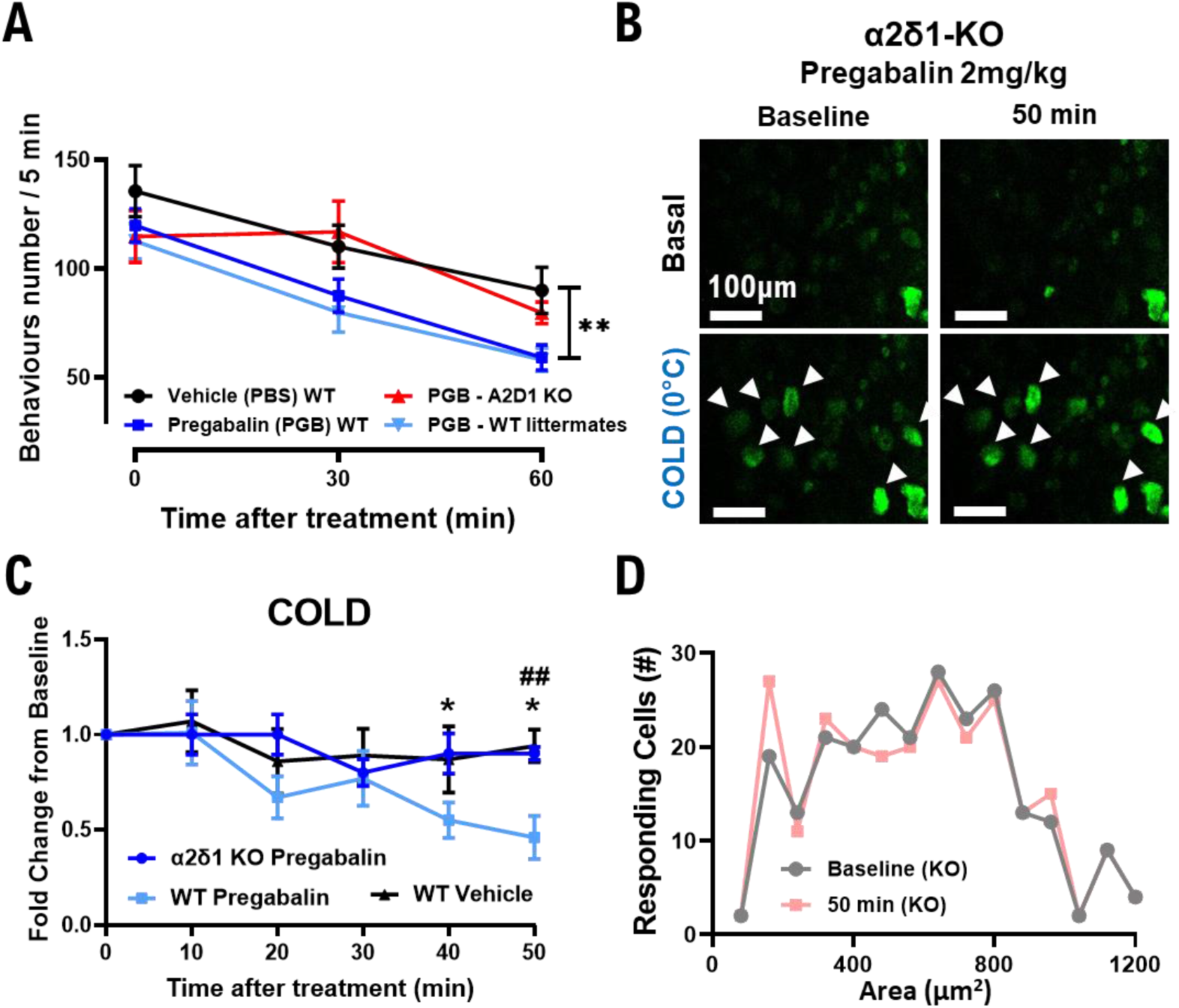
Knock-out of the α_2_δ-1 subunit of the VGCCs is sufficient to abolish the effects of pregabalin both on cold allodynia and on silencing of cold responding DRG neurons. (A) Plot showing the decrease of the number of nociceptive behaviours over time from vehicle (*n*=8) (black trace) or pregabalin injection in WT (*n*=8 (blue trace) + 12 (light blue trace)) and α_2_δ-1-KO mice (*n*=12) (red trace) when exposed to a 10°C cold plate for 5 minutes. (B) Example images showing that the reduction of the population of DRG neurons responding to ice-water 50 minutes after treatment with pregabalin is abolished in α_2_δ-1-KO mice. (C) Graph showing the disappearance of the decrease of the percentages of cell responding to ice-water stimulus in α_2_δ-1-KO mice. Change is quantified as fold-change to baseline. (D) Probability plots of the cell areas in oxaliplatin-treated α_2_δ-1-KO mice before (grey trace) and 50 minutes after pregabalin injection (pink trace). *n*=7 α_2_δ-1-KO mice, *n*=6 pregabalin-treated mice, *n*=5 vehicle-treated mice for imaging data. Statistical analyses in (C) was performed using Repeated Measures ANOVA test with multiple comparisons. * = P<0.05; ## = P<0.01 (α_2_δ-1-KO vs WT pregabalin-treated).

We then proceeded with *in vivo* imaging experiments on α_2_δ-1-KO mice. Similar to the results of the behavioural tests, the decrease of the percentage of cold-sensing neurons is completely abolished in the α_2_δ-1-KO mice, both in the case of ice-water and acetone stimuli (Figure 4B) (Supplementary Fig. 7A, B). This effect of α_2_δ-1 is specific again to cold stimuli, as the percentages of heat and mechanical-responding neurons did not vary significantly between conditions (Supplementary Fig. 7A, C, D). Furthermore, the cross-sectional area of the cold-responding neurons is similar to that of WT animals and did not vary upon treatment with pregabalin (Figure 4D) (Supplementary Fig. 8), consistent with the behavioural results and suggesting a strong dependence of the effect of pregabalin on its main molecular target α_2_δ-1.

## Discussion

### Pregabalin exerts a novel, peripheral effect on cold allodynia induced by oxaliplatin in mice

We have identified a novel effect of pregabalin on ‘silent’ cold sensing neurons that acquire de novo sensitivity to cold after cold allodynia induction by acute injection of oxaliplatin. Pregabalin, in this model, acts peripherally on this subpopulation of peptidergic A-fibre nociceptors by decreasing their excitability and thus mitigating the increased responses to cold stimuli that is thought to be a common pathophysiology underlying cold allodynia in various models of neuropathic pain^3^.

### Oxaliplatin mediates a *de novo* sensitization of previously silent nociceptors to cold stimulus: pregabalin reverts it

Chemotherapy-Induced Peripheral Neuropathy (CIPN) is a side effect of oxaliplatin treatment, as it is for many anticancer therapies. These drugs are alkylating agents that act through the formation of DNA-platinum adducts to cause DNA damage^15,20^. Patients treated with platinum-based drugs usually develop acute allodynia as well as chronic neuropathic symptoms like numbness, ataxia, loss of reflexes, and of course pain: these side effects are often time-and dose-related, and in the case of oxaliplatin they affect almost 90% of patients that undergo chemotherapy^2,15,21^.

Oxaliplatin-dependent neurotoxicity may develop in two distinct moments: an acute and reversible sensory neurotoxicity with cold-triggered paraesthesia and dysesthesia of the extremities that occurs within hours from the infusion, and a chronic sensory neuropathy, again with a stocking-glove distribution, that occurs after accumulation of the drug due to repeated infusions^15,20,22^. Many animal models of oxaliplatin-dependent neuropathy have been developed that replicate the acute and chronic symptoms experienced by human patients^4,23,24^: we chose a model of acute neuropathy to focus the attention on the early changes in cold perception and understand the molecular basis of cold allodynia. Indeed, there is strong evidence of changes in the coding of the responses to cold stimuli in the DRG: in the healthy state, only a limited population of small-diameter neurons responds to a cooling stimulus in a very specific fashion. After treatment with oxaliplatin, cold stimuli elicit the response of a population of previously silent neurons responding to both cold and noxious mechanical stimuli. The investigation of their molecular identity suggests they could be classified as A-fibre peptidergic nociceptors that express the sodium channel NaV1.8, whose key role in the sensation of prolonged, extreme cold in normal conditions has been previously shown^3,19^.

Pregabalin is a well-known anticonvulsant and analgesic and has been one of the staples in the treatment of neuropathic pain for years. Its use, alone or in combination with other drugs, has been shown to be effective in numerous preclinical and clinical models of neuropathic pain^14^. This drug has been shown to be particularly effective in the mitigation of cold allodynia, with single dose reversal of cold symptoms occurring after Spinal Nerve Ligation (SNL) and Spared-Nerve Injury (SNI) models as well as an attenuation of allodynia induced by oxaliplatin that showed pregabalin as the most efficacious drug alongside lidocaine and morphine as notable examples^9,14,23,25^.

This article confirms previous data showing the effectiveness of pregabalin in treating oxaliplatin-dependent symptoms and highlights its potential role as a functional counter to the increased excitability caused by the chemotherapeutic treatment^4,23,26^. The injection of 2 mg/kg of pregabalin ameliorated cold allodynia symptoms with a significant decrease in the number of nocifensive responses of mice exposed to a 10°C cold plate assay. Furthermore, the treatment caused a significant decrease of the number of neurons responding to different cold stimuli within the DRG to almost normal levels, with a time frame that corresponds to the effect seen on the behavioural tests and is in concert with other experiments done in similar conditions^23,27^.

Furthermore, there seems to be a direct effect on cellular excitability: the temperature threshold of cells responding to cold, that is not affected by oxaliplatin, shifts significantly towards lower temperatures after treatment with pregabalin, suggesting that pregabalin may silence cold-sensing neurons by increasing their response threshold.

Finally, the investigation on the changes in polymodality and cross-sectional area of neurons responding to cold before and after pregabalin injection strongly indicates that, albeit not being a direct counter to the alterations elicited by oxaliplatin, pregabalin indeed inhibits preferentially the mechano-cold ‘silent’ sensors that are the functional result of its pathological action.

Taken together, these data suggest that pregabalin acts directly on DRG cells by reverting the functional representation of cold stimuli back to almost normal levels, and this in turn ameliorates the symptoms of cold allodynia.

### Pregabalin efficacy is dependent on the α_2_δ-1 subunit of calcium channels

The main analgesic effect of pregabalin is assumed to be due to its action on the VGCCs through its direct binding with the α_2_δ-1 subunit. This subunit is expressed in primary afferent fibres and is fundamental for a correct release of neurotransmitters and neuropeptides at the presynaptic site in the dorsal horn of the spinal cord^16,28,29^. This protein has been extensively studied for its impact in neuropathic pain: it has been demonstrated to be upregulated in various models of neuropathic pain, and its upregulation alone is sufficient to drive allodynia, even in absence of a neuropathic injury^12,16,30^. Furthermore, it was observed that knockdown of the α_2_δ-1 subunit has an impact on acute sensation: α_2_δ-1-KO mice in fact have impaired mechanical and cold sensitivity, as well as showing a marked delay in the onset of mechanical hypersensitivity in a neuropathic model of Partial Sciatic Nerve Ligation (PSNL). Moreover, in the same model, the absence of α_2_δ-1 abolishes the anti-allodynic effect of a normally effective dose of pregabalin^18^.

We do not observe any effect of the α_2_δ-1-KO on the basal cold responses after oxaliplatin treatment, but we do show that the rapid effect of pregabalin on both the behavioural and functional representation of cold allodynia that were observed are almost completely abolished in the α_2_δ-1-KO mice. Indeed, the same treatment that elicited a significant reduction both in the number of nocifensive behaviours and of cold-responding cells did not have any effect in the α_2_δ-1-KO mice, further confirming the dependence of pregabalin on the α_2_δ-1 molecule for its early action in this CIPN pain model.

### Site of action of pregabalin: central, peripheral, or a bit of both?

Although gabapentinoids have been widely used in curing patients with neuropathic symptoms, their mechanism of action is not fully understood. They are thought to act mainly by interfering with trafficking of CaV2.2 VGCCs to the presynaptic site of the dorsal laminae in the spinal cord, as well as blocking their function through the α_2_δ-1 subunit, decreasing in turn the release of neurotransmitters and neuropeptides from DRG presynaptic sites and overall neuronal excitability in the Dorsal Horn (DH)^12,14^. However, decreased expression of VGCCs in the terminal may not be important, as their blockade by Mn^2+^ has a negligible effect on neurotransmitter release^16,31^. Other putative mechanisms of action include an inhibition of descending serotoninergic facilitatory and an excitation of descending noradrenergic inhibitory pathways, as well as an effect on synaptogenesis by blocking the interactions of thrombospondin to α_2_δ-1^32–34^. Another recent effect of gabapentinoids is the decrease of trafficking of NMDA glutamatergic receptors to presynaptic sites in the DH through a direct interaction of the C-terminus of α_2_δ-1 with these receptors^35^. Taken together, these findings show how pregabalin has many different targets and sites of action throughout the nervous system, corroborated by the recent discoveries of new interactions of the α_2_δ-1 protein, and by the fact that an increase of α_2_δ-1 expression has been observed in multiple sites within the peripheral and central nervous system^13,24,36,37^.

The overall site of action of pregabalin has been thought to be central, mainly at the spinal cord level but without ruling out the possibility of supra-spinal actions^12–14^. Indeed, electrophysiological studies on intact and spinalized rat showed that the inhibitory effect observed with pregabalin treatment on C-fibre-mediated nociceptive activity is abolished in spinalized animals, suggesting a strong involvement of supra-spinal centres^38^. Nevertheless, our data strongly suggest a direct effect on the excitability of a specific sub-population of peripheral DRG neurons, that acquire cold sensitivity after oxaliplatin-dependent neuropathic injury. Albeit we cannot infer a direct causal effect between the antiallodynic effect of pregabalin and the decrease in the population of cold-sensing cells at the DRG level, we suggest a strong correlation between the behavioural data and the functional changes in the peripheral representation of noxious cold stimuli. Indeed, it has been observed that gabapentinoids have a direct effect on the activity of specific DRG neuronal populations: gabapentin inhibits persistent sodium currents in medium sized neurons, and has an inhibitory effect exerted preferentially on small IB4-negative and medium-sized neurons^31,39^. The subpopulation of neurons preferentially hit by pregabalin, as shown in this paper, seems to be the silent cold sensors described previously^3^: they are medium-sized neurons, positive for NaV1.8 and CGRP. These results are further supported by microarray data on DRG of NaV1.8-DTA mice that show a strong reduction in the transcripts for CGRP-α and CGRP-β, as well as in CaV2.2 and α_2_δ-1^40^. Furthermore, it has been shown that gabapentinoids influence neurons of the substantia gelatinosa in the DH, with an overall decrease in excitability that cannot be explained just by a decrease of calcium currents, since, as was said previously, their blockade by Mn^2+^ does not affect overall neurotransmitter release in the DH^31^. On the other hand, IB4-negative neurons and Aδ fibres from medium-sized neurons project primarily to excitatory DH neurons^41,42^: if these cells are preferentially hit by gabapentinoids, we have also shown that the central action of pregabalin may in part depend on its selective inhibition of a specific subpopulation of silent cold neurons in the DRG.

## Conclusions

Overall, our results clearly indicate the presence of an acute, peripheral effect of pregabalin on cold allodynia induced in a CIPN model that is dependent on its main therapeutic target α_2_δ-1. This effect balances the peripheral sensitization observed via *in vivo* calcium imaging by decreasing the excitability and therefore silencing the medium-sized, A-fibre ‘silent’ cold sensors that become active in the DRG following oxaliplatin treatment. The imaging data collected here show the importance of having a functional representation of peripheral coding of pain and somatosensation in concert with behavioural data to better understand the changes elicited by pain models as well as visualize the effect of therapeutic drugs. Furthermore, they clearly provide new insights in the debate about a central versus peripheral effect of pregabalin, showing for the first time a peripheral effect of pregabalin on specific populations of DRG neurons *in vivo*.

## Acknowledgements

We thank Xinzhong Dong for the Pirt-GCaMP-3 mice line and Annette Dolphin for the α_2_δ-1-KO mice line.

## Figures

**Supplementary Fig 1.**
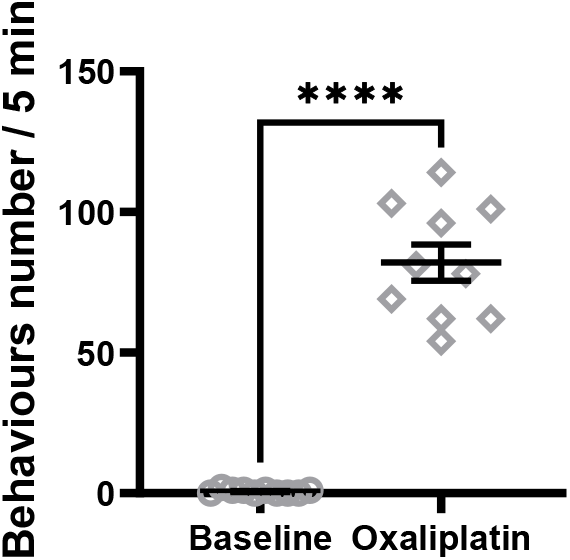
Assessment of cold allodynia after injection of oxaliplatin. Cold-plate assessment at 10°C of mice treated with oxaliplatin. Activity was measured as the total number of nociceptive behaviours (shaking, lifting, licking, guarding, biting) over the test duration. A cutoff time of 300s was used to limit tissue damage. *n* = 10 oxaliplatin-treated mice. Statistical Analysis was performed using paired t-test. **** = P<0.0001.

**Supplementary Fig. 2:**
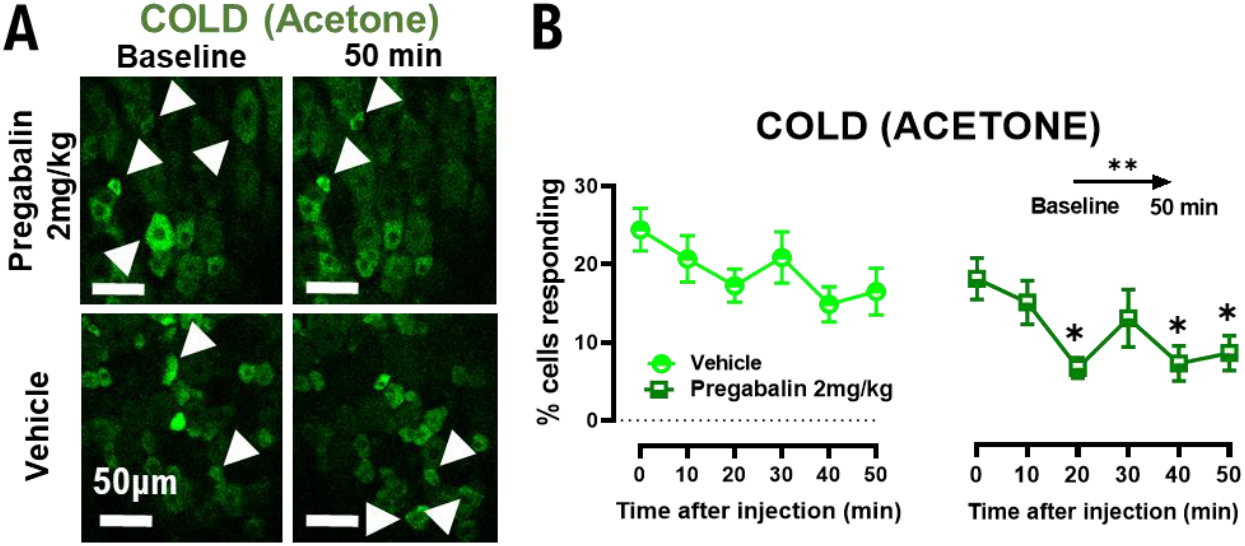
Pregabalin treatment decreases the number of neurons responding to a chemical cold stimulus. (A) Example images showing the reduction in the population of DRG neurons responding to acetone 50 minutes after treatment with 2 mg/kg of pregabalin. (B) Graph showing the decrease of the percentages of cell responding to acetone. *n* = 6 pregabalin-treated mice, *n* = 5 vehicle-treated mice. Statistical analysis in (B) was performed using Repeated Measures ANOVA test with multiple comparisons. * = P<0.05; ** = P<0.01.

**Supplementary Fig. 3:**
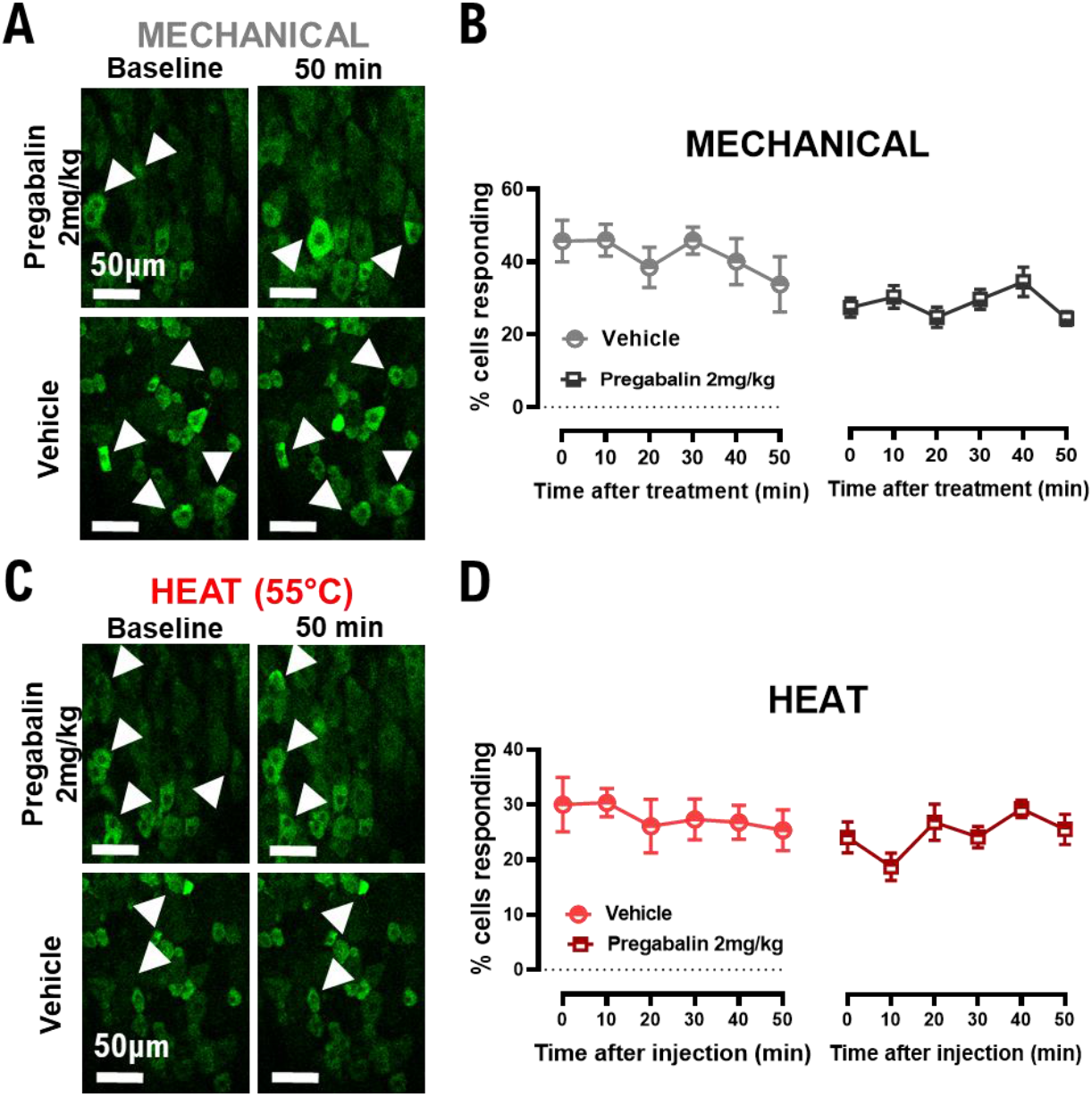
Pregabalin treatment does not affect other sensory modalities besides cold. (A) Examples images showing no change in the population of DRG neurons responding to mechanical pinch 50 minutes after treatment with 2 mg/kg of pregabalin. (B) Graph showing the unchanged percentages of cell responding to mechanical pinch. (C) Example images showing no change in the population of DRG neurons responding to a 55°C water stimulus 50 minutes after treatment with 2 mg/kg of pregabalin. (D) Graph showing the unchanged percentages of cell responding to a 55°C water stimulus. *n* = 6 pregabalin-treated mice, *n* = 5 vehicle-treated mice. Statistical analysis in (B) and (D) was performed using Repeated Measures ANOVA test with multiple comparisons.

**Supplementary Fig. 4:**
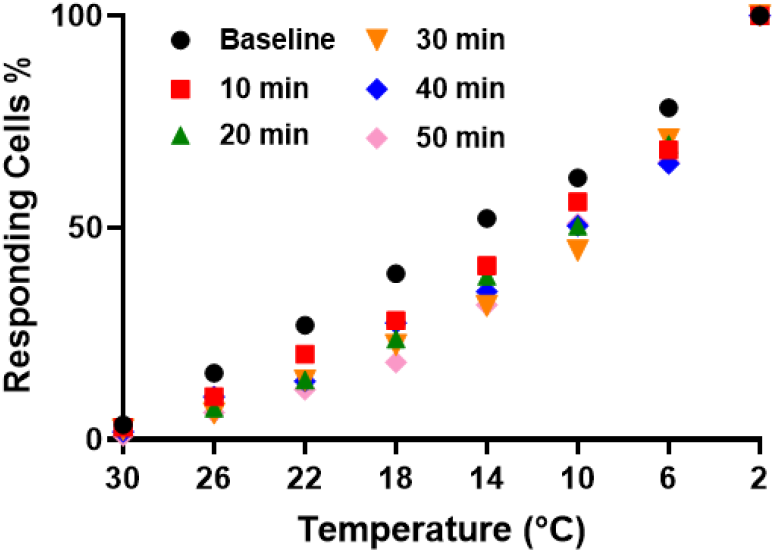
Pregabalin treatment increases temperature threshold of cold-responding cells over time. Graph showing the change of the relationship between the number of cold-responding neurons and the temperature drop over time from pregabalin injection.

**Supplementary Fig. 5:**
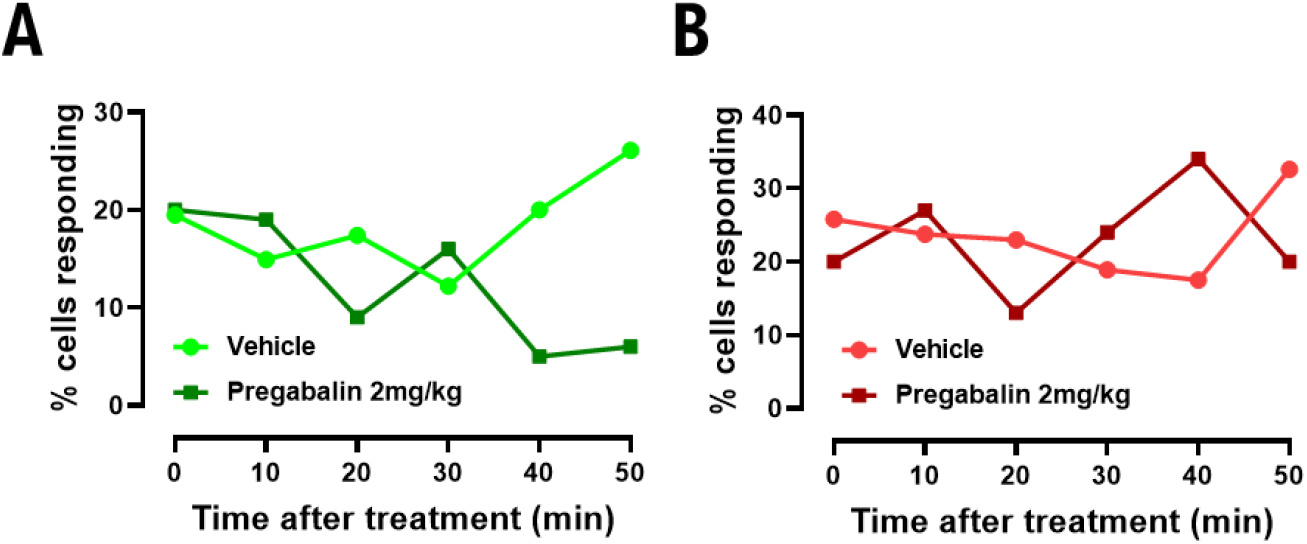
Pregabalin treatment reduces the number of polymodal mechano-cold but not mechano-heat responding cells. (A) Graph showing the change in the number of total mechano-responding neurons that respond also to acetone stimulus. Mice treated with vehicle do not exhibit the decrease over time from treatment as the mice treated with pregabalin do. (B) Graph showing the change in the number of total mechano-responding neurons that respond also to a 55°C water stimulus. Mice do not exhibit significant difference in the percentage of polymodal mechano-heat responding neurons over time from pregabalin treatment. *n* = 6 pregabalin-treated mice, *n* = 5 vehicle-treated mice.

**Supplementary Fig. 6:**
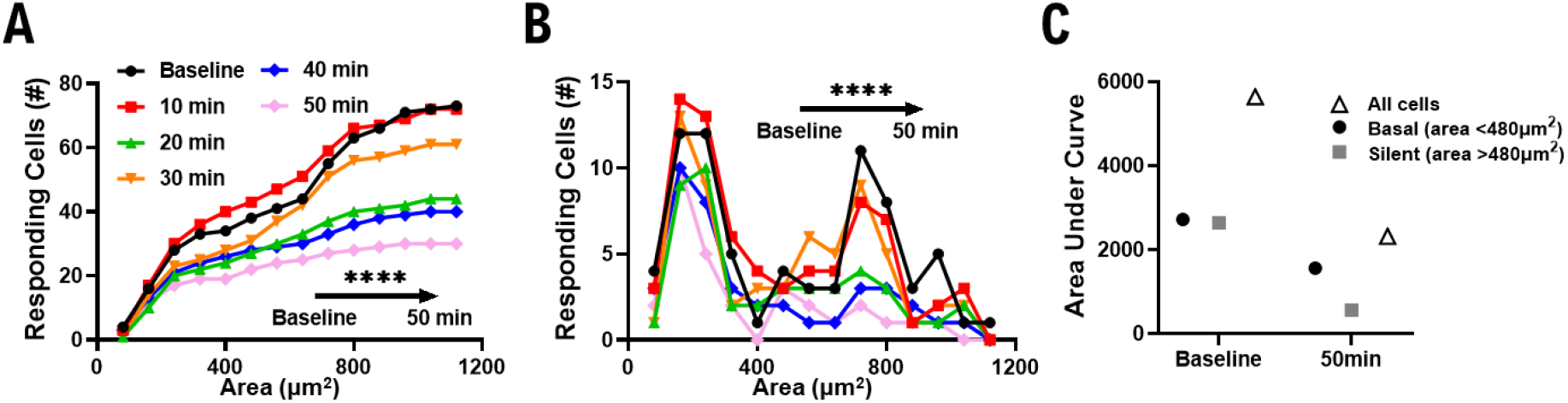
Pregabalin preferentially inhibits the activity of ‘silent’ cold-sensing neurons. (A) Cumulative plots of the cell areas in oxaliplatin-treated mice before and after pregabalin injection. The difference between the distributions at different timepoints after treatment is statistically significant (P<0.0001), as analyzed by Kruskall-Wallis test. (B) Number plots of the distribution of cell areas of cold-responding cells in oxaliplatin-treated mice before and after pregabalin injection. The difference between the numeric distributions at different timepoints after treatment is statistically significant (P<0.0001), as analyzed by Kruskall-Wallis test. (C) Graph showing the area under curve changes before and 50 minutes after pregabalin injection. The cell areas are divided into basal cold responding cells (A<480μm^2^) and silent cold sensors (A>480μm^2^). Albeit there is a difference in both populations after 50 minutes from treatment, the population of silent cold sensors seems to be silenced almost completely. These thresholds have been calculated in MacDonald et.al^3^.

**Supplementary Fig. 7:**
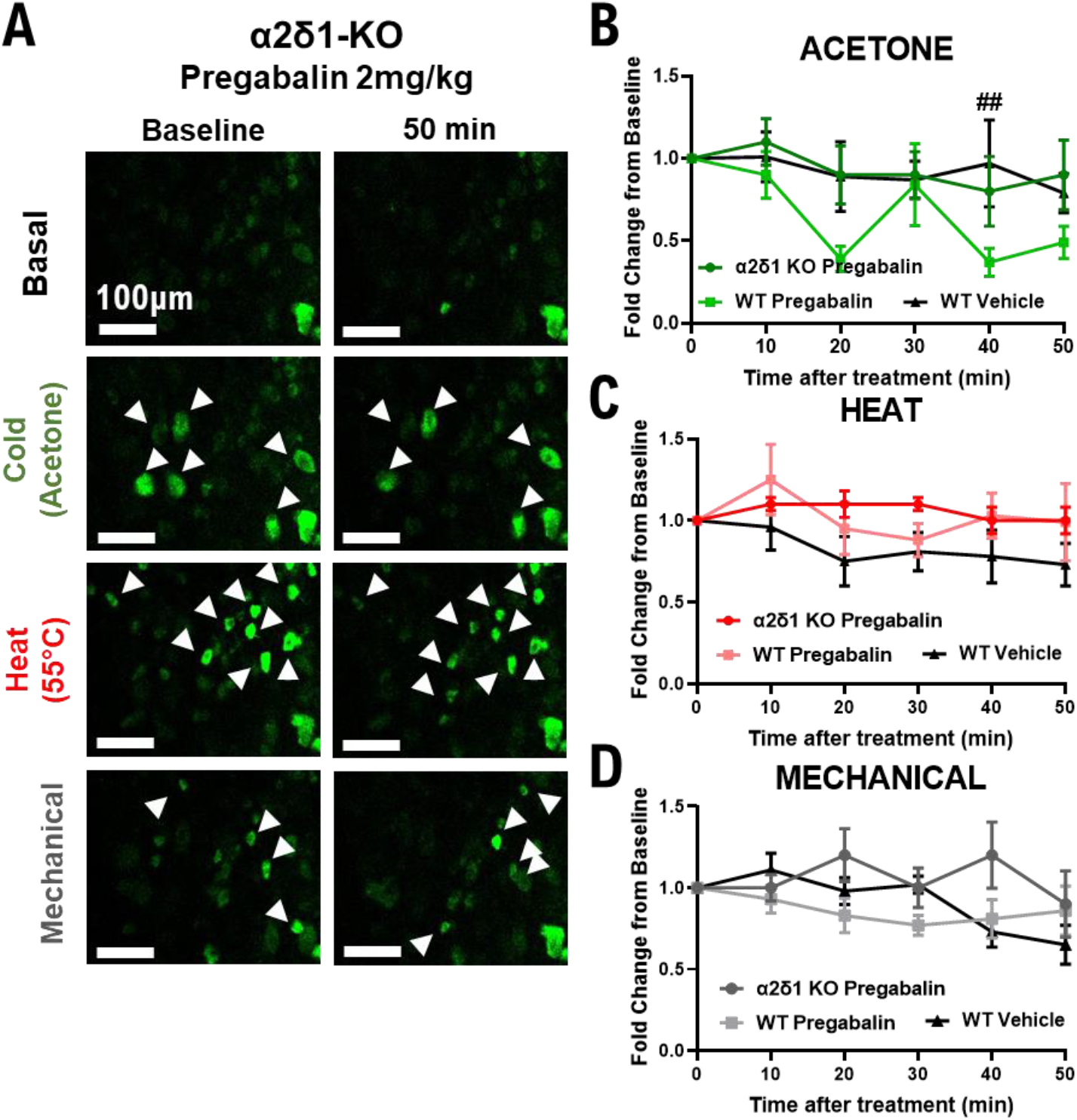
Knock-out of the α_2_δ-1 subunit of the VGCCs abolishes the effects of pregabalin on cold but has no effect on heat and mechanical responses. (A) Example images showing no change in the population of DRG neurons responding to acetone, 55°C water, and mechanical pinch 50 minutes after treatment of α_2_δ-1-KO mice with 2 mg/kg of pregabalin. (B) Graph showing the unchanged percentages of cell responding to acetone in α_2_δ-1-KO mice with respect to WT ones. (C) Graph showing the unchanged percentages of cell responding to a 55°C water stimulus in α_2_δ-1-KO mice with respect to WT ones. (D) Graph showing the unchanged percentages of cell responding to mechanical pinch in α_2_δ-1-KO mice with respect to WT ones. *n*=7 α_2_δ-1-KO mice, *n*=6 pregabalin-treated mice, *n*=5 vehicle-treated mice for imaging data. Statistical analyses in (B), (C), (D) were performed using Repeated Measures ANOVA test with multiple comparisons. ## = P<0.01 (α_2_δ-1-KO vs WT pregabalin-treated).

**Supplementary Fig. 8:**
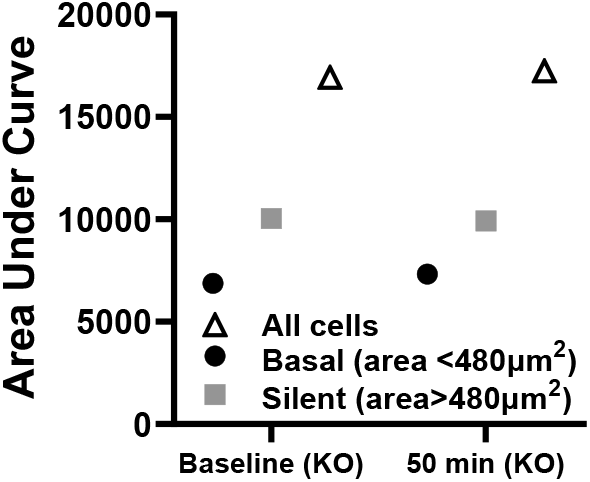
Knock-out of the α_2_δ-1 subunit of the VGCCs abolishes the silencing effect of pregabalin on silent cold sensors. Graph showing the area under curve changes before and 50 minutes after pregabalin injection in the α_2_δ-1-KO mice. The cell areas are divided into basal cold responding cells (A<480μm^2^) and silent cold sensors (A>480μm^2^). There seems to be no difference in the population of basal and silent cold sensors after pregabalin treatment when the α_2_δ-1subunit is knocked down globally. These thresholds have been calculated in MacDonald et.al^3^.

